# Precision measurements of brain oxygen utilization with positron emission tomography

**DOI:** 10.1101/2025.02.16.638562

**Authors:** John J. Lee, Joshua S. Shimony, Nicholas V. Metcalf, Andrei G. Vlassenko, Manu S. Goyal

## Abstract

Oxygen utilization is important for studies of brain metabolism, alongside other measurements such as for glucose metabolism. Oxygen and other measurements with [^15^O] tracers and PET, however, are significantly more challenging than measurements of [^18^F]fluorodeoxyglucose, the standard for probing tissue glucose metabolism in vivo, in part due to the much shorter radioactive half-life of [^15^O]. This work examines details of precision measurement of [^15^O] tracers and their kinetics. Investigations of arterial input functions (AIFs) and image-derived input functions (IDIFs) have figured prominently for PET, but [^15^O] tracers are rarely studied given the small numbers of PET facilities equipped to work with these tracers, particularly in their inhaled form. Estimates of IDIFs and AIFs for [^15^O] tracers have demonstrably distinct characteristics arising from instrumentation as well as circulatory physiology. To reconcile IDIFs and AIFs, we developed a generalizable model for bolus tracer transport, corrected for known effects of instrumentation for measuring IDIFs and AIFs, and found intravascular [^15^O]CO to be especially suited for constructing a robust recovery coefficient for IDIFs compared against AIFs. Within a Bayesian framework for posterior estimation and estimating data evidence, IDIFs provide parameter estimates compatible with AIFs in the setting of biological variability. IDIFs also provide data evidence that exceeds that of results from AIFs. These suggest that scalar recovery coefficients may be adequate to estimate partial volume effects, and that the circulatory consistency of internal carotid IDIFs with brain tissue perfusion provides greater precision than what can be estimated using radial artery AIFs, which exhibit greater variability of recirculation wave forms.

## Introduction

Positron emission tomography (PET) of oxygen utilization has importance for foundational studies of brain activity. In the presence of physiological neural activity (e.g., visual stimulation by annular reversing checkerboard), oxygen metabolism rises modestly compared to glucose metabolism and blood flow [1], a disassociation which revealed the importance of metabolic mechanisms beyond energy metabolism [1]–[3]. Furthermore, regional distributions of aerobic glycolysis [3], which require measurements of both glucose and oxygen metabolism, are associated topographically with development, brain aging, and neurodegeneration [4]–[8].

However, the most challenging measurements of metabolism in the brain involve oxygen utilization [9]. [^15^O] emits purely positrons, without prompt gammas [10]. Nevertheless, positrons from [^15^O] have significantly greater free paths of travel in tissue, and correspondingly greater annihilation energies, greater than all other common pure-positron emitting radioisotopes [10]. These longer free path ranges increase partial volume effects. Furthermore, high-energy annihilation gammas produce Compton scattering and pair production of *e*^+^*/e^−^*, further degrading spatial resolution. High-energy positrons also escape human skin, travel through air, and annihilate at the scanner’s internal tunnel, affecting image quality.

Previous studies of the brain with older PET scanners required AIFs due to their poorer quality dynamic data with limited spatial resolutions [11]. Modern scanners are markedly improved and radial artery cannulation safety considerations have evolved, given the increased importance of the radial artery for modern medical interventions. These evolutions of the field further the merit of developing precision quantitative PET using noninvasive methods. However, IDIFs have demonstrated differences in physiologic metrics, compared to AIFs, that produce unrealistic physiological metrics, and these have been mostly attributed to partial volume effects. In previous studies, rescaling has appeared effective, with rescaled IDIFs providing reduced intersubject variability and challenging whether AIF analysis should be considered as a gold standard [12].

This raises questions of model selection and data selection. Dynamic nested sampling, a Bayesian technique, can provide helpful metrics for identifying models and data that are mutually consistent, and thereby increase the precision of PET measurements. In this work, we develop an IDIF model with dynamic nested sampling to show that [^15^O] PET imaging can be quantified non-invasively and comparably to invasive measurements. We first describe our operational procedures and a novel phenomenological model for input functions. We introduce the advantages of dynamic nested sampling for model selection. Next, using detailed exemplars drawn from our results, we illustrate key features of circulatory hemodynamics that influence AIFs and IDIFs. We illustrate how intravascular tracers such as [^15^O]CO may be helpful for estimating recovery coefficients that reconcile AIFs with IDIFs. Finally, we systematically examine how AIFs and IDIFs influence topographically organized estimations of physiologically pertinent metrics of hemodynamics and oxygen metabolism.

## Materials and Methods

### Study participants

All human participation contributing to this study conformed to the Declaration of Helsinki, including consenting to participation, and oversight by the Washington University School of Medicine Institutional Review Board and the Radioactive Drug Research Committee. Inclusion criteria were expansive, but limited to adults, and exclusion criteria were limited so as to obtain generalizable results from six participants. Exclusions included neurological, psychiatric or general medical conditions that might preclude their ability to complete the study. Specific exclusions also included contraindications for MRI or PET scanning, pregnancy, and medication use that could significantly influence study measurements. These participants consenting to invasive arterial procedures were drawn from larger on-going studies of non-invasive studies of brain metabolism and aging [8].

### MRI and PET scanning with [**^15^**O]CO, [**^15^**O]O**_2_**, and [**^15^**O]H**_2_**O

MRI included 3D magnetization-prepared rapid gradient echo (MPRAGE) pulse sequence acquisitions with 0.8 mm isotropic voxels and time-of-flight magnetic resonance angiography (MRA) reconstructed to 0.26 × 0.26 × 0.5 mm^3^ voxels from 3T Siemens Prisma scanners. FreeSurfer version 7.3.2 [13], [14] produced segmentations combined with cortical parcellations from the Schaefer 200 atlas [15]. All PET scanning ensured that participants remained awake with eyes closed. Head movements were minimized by elastic head restraints. PET scans were acquired on a single Siemens Vision 600 scanner equipped for contemporaneous CT scanning for construction of attenuation maps. Dosing for [^15^O]CO and [^15^O]O_2_ ranged 10-46 mCi. Dosing for [^15^O]H_2_O ranged 21-27 mCi. Participants received training for inhalation of gaseous tracers through polypropylene breathing tubes prior to scanning. Comprehensive details of these PET scanning protocols have been reported [16].

### Invasive arterial sampling

Arterial blood measurements rely on expert cannulation of the radial artery. We relied on a centralized arterial line service at the Center for Clinical Imaging Research at Washington University School of Medicine, staffed by attending interventional physicians and clinical fellows working within their scope of practice in either interventional radiology or anesthesiology. Following cannulation, nursing staff maintained pressurized catheter line sets, ensuring clinical safety, and operating valves, pumps, and drains during repeated blood measurements. Nurses drew arterial blood through a line circuit comprising serial sets of extension catheters, multi-way valves, and a compact, tungsten-shielded LYSO coincidence detection probe (Twilite II, swisstrace.com), at 5 mL/min by peristaltic pump (MRidium, iradimed.com). Scintillation photons from the probe head transmitted along light guides to photomultipliers and timing electronics, following which data acquisition software recorded coincidence counts with rated efficiency of 2.4 cps/kBq/mL. We previously reported comprehensive details of the materials, methods, and protocols employed in collecting these data [16].

### Emissions scanning and reconstructions from listmode

We obtained static and dynamic reconstructions of emissions at the scanner console for quality assurance. For all quantitative measurements, we used listmode emissions data for computationally intensive reconstructions and analyses. This report’s software archive includes classes and unit tests in Matlab that describe high-level implementations of JSRecon12, e7 tools and Brain Motion Correction (BMC) (https://github.com/jjleewustledu/mlsiemens). We implemented software components with versions VG80 for e7, 27-Sep-2023 for BMC, 14-Jul-2023 for JSRecon12. Listmode data for participant 1 required version VG76 for e7, consistent with the scanner’s operating environment at the time of scanning. Essential choices for configuring our reconstructions included: time-of-flight reconstructions with point-spread function corrections, 4 iterations and 5 subsets for ordinary-Poisson ordered sets of expectation maximization, absolute scatter correction, 440 × 440 matrix size, and no postfiltering. To improve signal-to-noise, but preserve adequate spatiotemporal resolution for generating IDIFs matched to sampling characteristics of invasive AIFs, we constructed moving windows, using 10 s frame durations and moved the start of frames by 0-9 s, thereby generating 290 interleaving frames for 299 s of listmode [^15^O]CO and 120 interleaving frames for 129 s of listmode of [^15^O]H_2_O or [^15^O]O_2_. Some acquisitions contained longer durations of listmode, which we truncated to 299 s and 129 s, respectively. BMC infrequently failed to reconstruct imaging frames late in the scanning of [^15^O] tracers, which we confirmed to exhibit marginal signal-to-noise characteristics in quality assurance data, and these empty frames were omitted from further inference and analyses.

### Image-derived input functions from internal carotid artery centerlines

IDIF constructions used centerlines traversing the centers of the petrous segments of the internal carotid arteries. These constructions used co-registered magnetic resonance angiography (MRA), magnetization-prepared rapid gradient echo (MPRAGE), and static PET emissions reconstructions, from which we generated maximum intensity projections along *x, y*, and *z* axes, then manually traced centerlines consistent with all imaging modalities. While less scalable, manually drawn centerlines most closely conform with the visualized anatomy, and we chose this workflow as a gold standard. Notably, clinical radiology workflows commonly provide automations for generating centerlines.

Figure 1 provides comprehensive views of arterial blood measurements for administration of [^15^O]CO, [^15^O]O_2_, and [^15^O]H_2_O. The origin of the time axis denotes the start of emissions scanning. Inhalation of gaseous [^15^O]CO, [^15^O]O_2_ and intravenous injection of [^15^O]H_2_O commenced simultaneously with commencement of scanning, except for [^15^O]CO for study participant 1, in which case scanning commenced approximately 80 s after inhalation of tracer. Loss of timing precision involved variations in inhalation behaviors of participants, although inhalation procedures were prescribed and practiced in advance of scanning. While personnel administrating intravenous injections meticulously timed injections, injection rates necessarily varied according to the compliance and capacity of the cannulated vein. Additionally, the venous flow to the heart has greater biological variability than that of pulmonary veins. Consequently, the delay of emergence of the tracer bolus in arterial blood measurements varied widely, from less than 20 s for inhaled tracers to greater than 40 s for intravenous tracer. The dispersion of arterial blood measurements is sensitive to the path of the tracer through biological compartments. The least dispersion is apparent for [^15^O]CO, which is directly inhaled, has no systemic venous paths prior to bolus measurement, and is intravascular, bound in excess of 98 % to formed elements of the blood in direct assays of whole blood and plasma(supplemental appendix 3). Dispersion was greater for [^15^O]O_2_, which is also directly inhaled without systemic venous paths, but is not intravascular, and may experience non-negligible uptake by the myocardium and arterial endothelium. Additionally, arterial blood measurements of [^15^O]O_2_ demonstrated bimodal bolus peaks or prolonged bolus peaks, suggesting the role of more variable respiratory inhalation. Dispersion was intermediate for [^15^O]H_2_O, which experiences protracted systemic venous paths and is diffusible across the myocardium and vascular endothelium.

**Figure 1.**
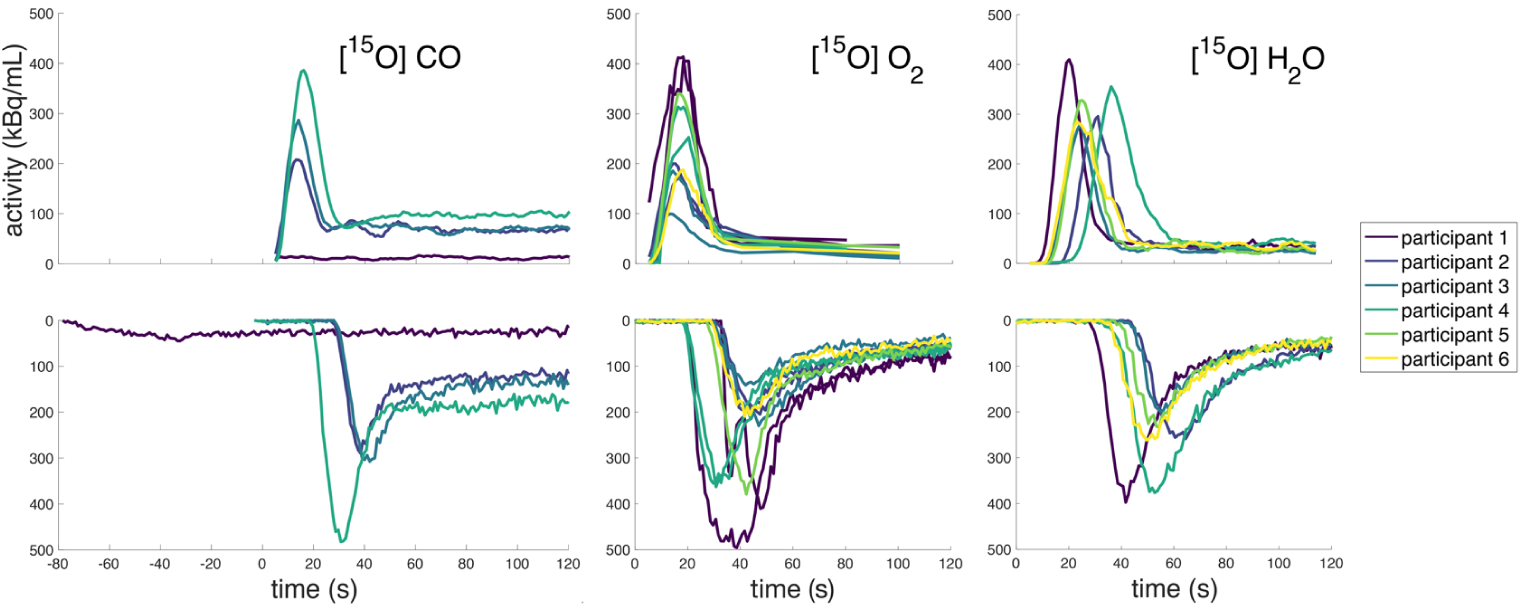
Uncorrected input functions for [^15^O] tracers. The **top row** illustrates IDIFs obtained by sampling centerlines from MRI in dynamic emissions reconstructions, moving 10 s frames by 1 s intervals in the mid-frame time. The origin of the time axis corresponds to the first mid-frame time. The **bottom row** illustrates AIFs obtained from arterial blood measurements with automated extraction of blood from the radial artery and continuous coincidence counting at a detection probehead. While both [^15^O]CO and [^15^O]O_2_ are inhaled gaseous tracers, the intravascular binding of [^15^O]CO dampens the shape of the bolus, providing similarity of bolus shapes across persons. For completeness, [^15^O]CO for participant 1 is illustrated despite premature inhalation of tracer before scanning and low activity of this participant’s dosing. Comparatively, [^15^O]O_2_ demonstrates significant variability across participants and also for test-retest repetitions of [^15^O]O_2_ scanning for participants 1-4. Bolus shape variations may be due to irregularities of forced inspiration of gases, possibly including multiple breaths. [^15^O]O_2_ also has avid uptake by tissues during recirculation, adding complex variations. As an intravenous tracer, [^15^O]H_2_O has more reproducible bolus shapes, like [^15^O]CO.

### Phenomenology of input functions

Historically, studies of input functions for blood circulations have discussed indicator dilutions in fluid models of continuity [17]–[19], commonly using phenomenologies based on the gamma distribution, which is the integrand of the Bernoulli gamma function [20], [21]. We extended a previously reported phenomenology of input functions, comprising gamma distributions, which had been constructed for the intravascular bolus-passage of gadolinium in dynamic susceptibility contrast MRI [22]. Our extensions generalize gamma distributions to feature a stretched exponential that provides a Kohlrausch-Williams-Watts shape parameter [23], [24]. Consistent with previous studies, our input functions describe a first passage of bolus, a single bolus describing early recirculation, and an asymptotic bolus describing highly dispersed late recirculations [22]. In this study of inhaled gaseous tracers, multiple inhalations and irregular inhalations can induce recirculation waveforms with significantly greater complexity than what has been historically described. In all, we articulate our phenomenological input function *ρ*_IF_ with

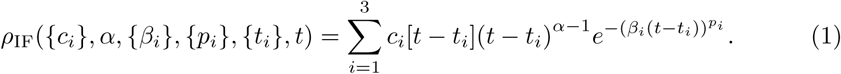

The first passage of the bolus is denoted with *i* = 1, and recirculations are denoted with *i >* 1. Scaling functions *c_i_*[*t* − *t_i_*] are also temporal step functions, zero for negative arguments in square brackets. The traditional shape parameter of the gamma distribution is *α*, describing a power law for temporal dispersions, and which we set identically for all bolus terms. The correspondingly traditional scaling parameter for temporal decays is *β_i_*, which we set identically for all bolus terms except the asymptotic term, *i* = 3, which describes persistent intravascular tracer experiencing exchange processes across the blood-brain barrier, while also experiencing slow elimination by hepatic or renal mechanisms. Fitted *β*_3_ were typically two orders of magnitude smaller than for non-asymptotic terms. By fixing *α* and *β* for non-asymptotic terms, we imposed an invariance of shape and scale that could be ascribed to the fixed geometry of fluid dynamics under vascular confinement. Assuming invariance of cardiac output during scan sessions was consistent with scanning conditions for all human study participants. The hypothesis of invariant parameters originated in previous work with intravascular gadolinium for MRI [22]. In numerical experiments, we could not identify an invariant Kohlrausch-Williams-Watts shape parameter, *p_i_*, for bolus terms, although imposing monotonic decrements, *p*_1_ *> p*_2_ *> p*_3_, appeared consistent with measurements. The Kohlrausch-Williams-Watts shape is suited for the dynamical complexity of paths traversed by tracer circulation and recirculations. As a final simplification, we identified *t*_3_ = *t*_1_, assuming the asymptotic bolus to have time of emergence identical to first passage of the first bolus term. We interpreted the second recirculating bolus, which traverses the entirety of the arterial-capillary-venous circuit, to have independent *t*_2_ ≠ *t*_1_.

Figure 2 illustrates the effects on variations of model parameters on the waveform of the unseen input function, which may only be observed with significant instrumental effects of delays, dispersions, and time averaging. For concreteness, all variations are perturbations of fitted model parameters for an AIF and an IDIF measured from [^15^O]O_2_ of study participant 1, the direct measurements of which are visible in figure 1. We note the presence of dual peaks of the input function in inferences of phenomenology corresponding to AIF and IDIF measurements. The variability of bolus arrival time *t*_0_ is especially consequential for AIF measurements, as they are necessarily significantly delayed by their long transit through the brachial and radial arteries, and 10 seconds of delay and dispersion through cathethers that cannulate the radial artery. Our phenomenology allows for temporal separation of dual peaks, *τ*_2_. The traditional shape parameter for gamma distributions is *α* − 1, and the traditional temporal scale parameter is *β*. However, these alone were inadequate for fitting our observations of input functions, leaving systematically visible residuals. The Kohlrausch-Williams-Watts parameters, *p*, *δp*_2_, and *δp*_3_, provided appropriate flexibility for fitting recirculation waveforms, which experience progressive dispersions with successive recirculations. The phenomenology usefully limits the asymptotic recirculation scale, 1*/γ* to the long-time waveform. The fractional contributions of the second peak, *f*_2_, and the asymptotic waveform, *f*_3_, provide linear superposition of waveform components.

**Figure 2.**
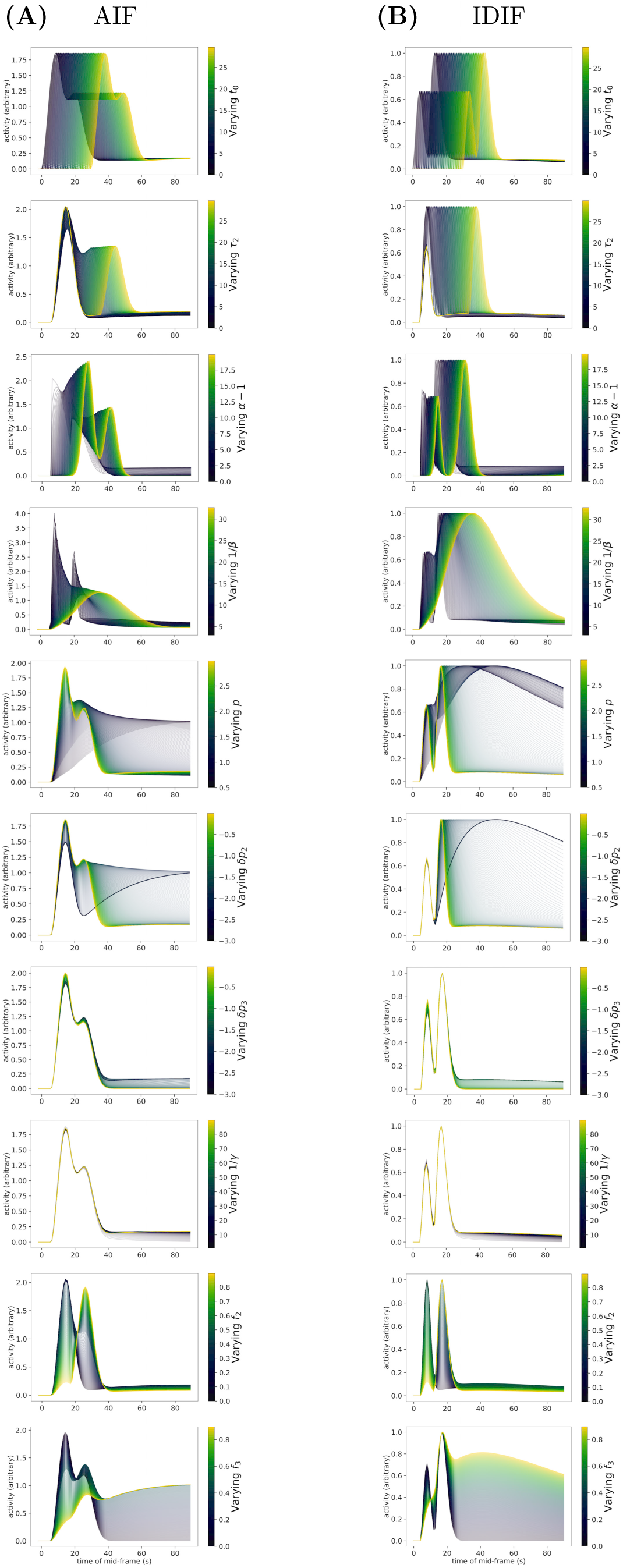
Variations of parameters for the model of input functions. We constructed a phenomenological model to represent the true activity of [^15^O] tracers in blood, *in vivo*. As detailed in methods, the model elaborates familiar formalisms based on series of generalized gamma distributions, which encode plausible physical features of a Kohlrausch-Williams-Watts function (non-equilibrium mechanics, linear relaxation, nonlinear energy dissipation). The model specifically facilitates use with numerical corrections of the influence of instrumental measurement procedures such as arterial blood sampling by arterial cannulation for arterial input functions and moving window reconstructions of listmode emissions for image-derived input functions. Subfigures illustrate variability of fitted AIF **(A)** and IDIF **(B)** by model parameters. Timing parameters for the intrinsic waveform of an input function are: arrival time *t*_0_ for the primary peak of bolus passage, delay *τ*_2_ for a secondary peak, and delay *τ*_3_ for a tertiary peak. Parameters for the empirical exponent of the stretched exponential are: *p* for the primary peak, *δp*_2_ for variations in the secondary, and *δp*_3_ for variations in the tertiary. Shape and scale parameters familiar for gamma distributions are: *α −* 1 for the power law in time, and 1*/β* for the exponential decay. 1*/γ* is the time constant for the asymptotic recirculation of blood. Coefficient weights for terms of the formal model series are: *f*_2_ weighting the secondary peak and *f*_3_ weighting the tertiary recirculation. Compared to equation 1, *τ*_2_ = *t*_2_ *− t*_1_; *p* = *p*_1_; *δp*_2_ = *p*_2_ *− p*_1_; *δp*_3_ = *p*_3_ *− p*_2_; 1*/γ* = 1*/β*_3_; *A*(1 *− f*_2_ *− f*_3_) = *|c*_1_|; *Af*_2_ = *|c*_2_|; *Af*_3_ = *|c*_3_|; and *Aσ* is the standard deviation of a Gaussian likelihood objective.

To reduce risks of overfitting and excessive researcher degrees of freedom, we performed nearly all training of phenomenological models using data from participant 1. Such training included selection of number of terms of the generalized gamma series, the prior ranges for parameters, and details of numerical implementation, such as convolutions, interpolations, and enforcements of numerical characteristics such as positivity (e.g., *ρ >* 0) or ordering of parameter magnitudes (e.g., *p*_1_ *> p*_2_ *> p*_3_). Initial estimates of prior ranges used values available from previous reports of [^15^O] PET [9], [25], [26]. Because of improved fits for participant 1 and the large capacity of models for dynamic nested sampling, we found it advantageous to allow adjustable values for physiological metrics such as the fraction of water of metabolism, the capillary blood volume, and post-capillary blood volume, which had previously been assume invariant [9]. Initial training on participant 1 used an expression for the asymptotic waveform ∼ (1 − exp −*β*_3_*t*), which had been previously reported [22]. However, following inference on all participants of this study, and noting the performance of our phenomenology on longer-lived radiotracers based as [^18^F], yet unreported, we adjusted our phenomenology such that the asymptotic waveform ∼ *t^α^* exp (−*β*_3_*t*)*^p^*^3^, which improved the agreement of predictions with measurements in a single, repeated round of inference.

### Dynamic nested sampling provided parameter estimations and data evidence

By construction, nested sampling algorithms directly estimate likelihood functions with respect to prior mass. Consequently, numerical integration immediately yields model evidence (marginal likelihood, marginal density of data, prior predictive, partition function) [27] from

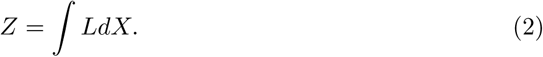

The likelihood *L* = *L*(*θ*) is a function of model parameters *θ*, and differential prior mass is *dX* = *π*(*θ*)*dθ*. In the context of Bayes’ theorem, the likelihood × prior = evidence × posterior. Viz.,

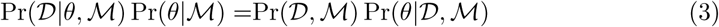

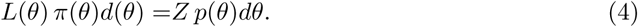

The observed data, D, modulates the prior, *π*(*θ*)*dθ*, for the posterior *p*(*θ*)*dθ*. The evidence, *Z*, encodes data conditional on model assumptions and has invariance (to estimation error).

We constructed models and inferences on our data implementing the dynesty framework [28]. Dynesty provides open-source implementations of dynamic nested sampling, featuring traditional Bayesian parameter estimation from posteriors as well as data evidence from posterior masses. Dynamic nested sampling provided inferential environments within which we constructed a hidden-variable model of the intrinsic input function that supplies brain tissues. We equipped the intrinsic input function to describe multiple boluses from inhalations, or, potentially, difficult venous injections, and to describe recirculation waveforms. We fitted the intrinsic input function with separate experimental measurements of the dispersion and delay of arterial blood measurements and the moving window averaging of our dynamic reconstructions of emission imaging. These allowed us to query our data for the intrinsic input function waveforms that were compatible with AIFs and IDIFs. These also allowed for calculations of data evidence for AIFs and IDIFs. This report’s software archive includes all python classes and jupyter notebooks, for quality assurance, which we used for nested sampling of our data (https://github.com/jjleewustledu/idif2024/tree/master).

### Instrumental cross-calibrations

We cross-calibrated all activity measurement instruments using aqueous solutions of [^18^F]FDG at approximately 50-100 kBq/mL. We mixed the calibration solution by shaking in a 650 mL polyethylene terephthalate media bottle (Nalgene^TM^, thermofisher.com), which we scanned with configurations for [^18^F]FDG PET. We also drew aliquots and measured activities in our arterial coincidence detection probe (Twilite II, swisstrace.com) as well as in our NaI well counter (Caprac, mirion.com). The well counter accommodated absolute calibration against sealed [^68^Ge] reference rod sources providing approximately 1 kBq activity (GF-068-R3, ezag.com). All measurements, other than for rod sources, were contemporaneous within the lifetime of the tracer. We adjusted all instrument efficiencies to match reference rods sources.

We measured the performance characteristics of our arterial catheter assembly in a single separate experiment. We mixed expired packed red cells diluted in plasma to several hematocrits, measured by centrifugation under clinical laboratory conditions. We split the reconstituted blood into approximate halves and mixed one half with [^18^F]FDG to approximately 50-100 kBq/mL. We supplied unlabeled blood to the catheter assembly, arterial coincidence detection probe, and peristaltic pump at target rate of 5 mL/min. We switched to labeled blood by manual switching (less than 0.5 s delays) to generate a Heaviside input to the arterial measurement instruments. These measurements provided hematocrit dependent impulse responses for the *ex vivo* catheter line assemblies used in arterial blood measurements.

The recovery coefficient was constructed from 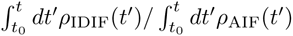, which included all overlapping times between arterial blood measurements and emissions scanning.

## Results

### Input functions require adjustments for sampling characteristics

Figures 3-6 illustrate exemplar results for dynamic nested sampling on [^15^O]O_2_ in study participant 1. Figure 3 highlights comparisons of two sequential administrations of [^15^O]O_2_, and the measurements of input functions by AIFs and IDIFs. The severe dispersion of AIFs substantially influences estimations of the amplitude of phenomenological input functions. Despite the contrasts between measurements, trace plots from dynamic nesting sampling demonstrate that most phenomenological parameters have meaningful posteriors with central tendencies. The lack of central tendency for *δp*_3_ and 1*/γ* is anticipated by their fit against long-time data samples for which [^15^O] has decayed substantially. The noninformative distributions for *A* demonstrate that inference of the input function amplitudes follow inferences on the more information distributions of other waveform parameters. Figure 4 verifies the weakly informative or noninformative distributions for *δp*_3_, 1*/γ*, and *A*, which have little covariance with other parameters. Run plots illustrate that the posteriors for IDIFs encompass greater volumes of the space of parameter configurations compared to AIFs. Dynamic nested sampling on physiological models, shown in figures 5 and 6, reveal similar trends. All physiological trace plots demonstrate central tendencies, and run plots reveal that IDIF parameters cover more volume of parameter configuration, suggesting that IDIF models may be more robust during inference.

**Figure 3.**
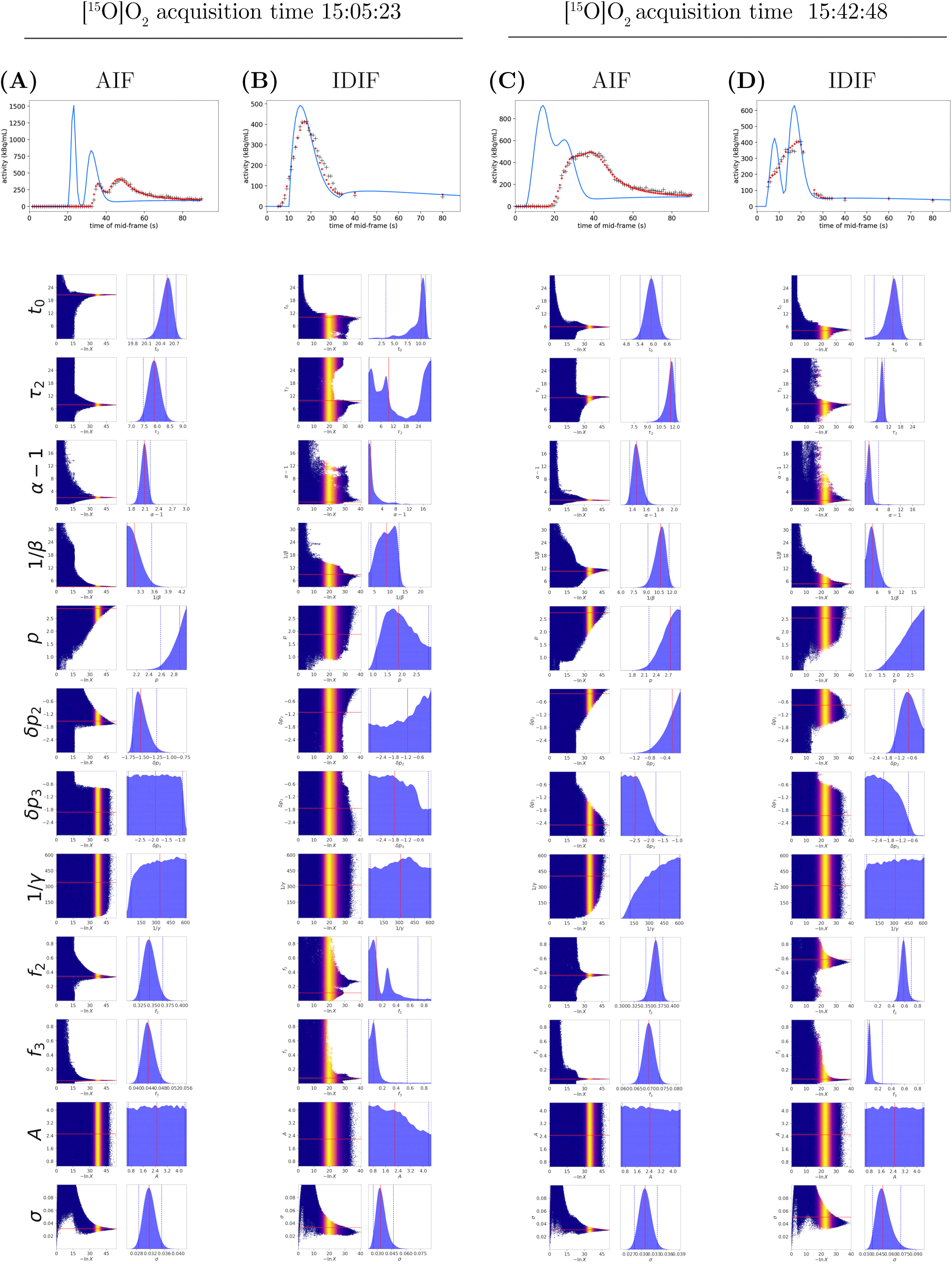
Dynamic nested sampling fits the bolus circulation model to repeated measurements of inhaled [^15^O]O_2_. Panels of column **(A)** illustrate an arterial input function (AIF), measurements of arterial blood sampled by cannulation of the radial artery, peristaltic pumping, and coincidence detection of annihilation photons within a compact detector probe. Panels of column **(B)** illustrate an image-derived input function (IDIF) sampled from centerlines through the petrous carotid artery, visible in dynamic PET tomography. Panels of columns **(C)** and **(D)** similarly illustrate an AIF and an IDIF obtained from a second dose of [^15^O]O_2_ which was administered 37 minutes following the first dosing. **Topmost panels** are *residual plots*. Blue curves depict the parametric model of unseen input functions, AIF or IDIF, while red dots show the transformation of unseen input functions to predictions of instrumental measurements, which involve delays and dispersions in AIFs and moving windows providing temporal averaging over dynamic PET tomography for IDIFs. Crosses represent measured data for AIFs and IDIFs. **Lower panels** are *trace plots* for model parameters. Vertically stacked dead points explore model parameter values in the course of a nested sampling run, depicted progressing left to right, with warmer colors indicating importance weighting of the probability density, *p*(*X*). Histograms of parameters values are medium blue, and the best fitted parameter values are marked by red vertical lines.

**Figure 4.**
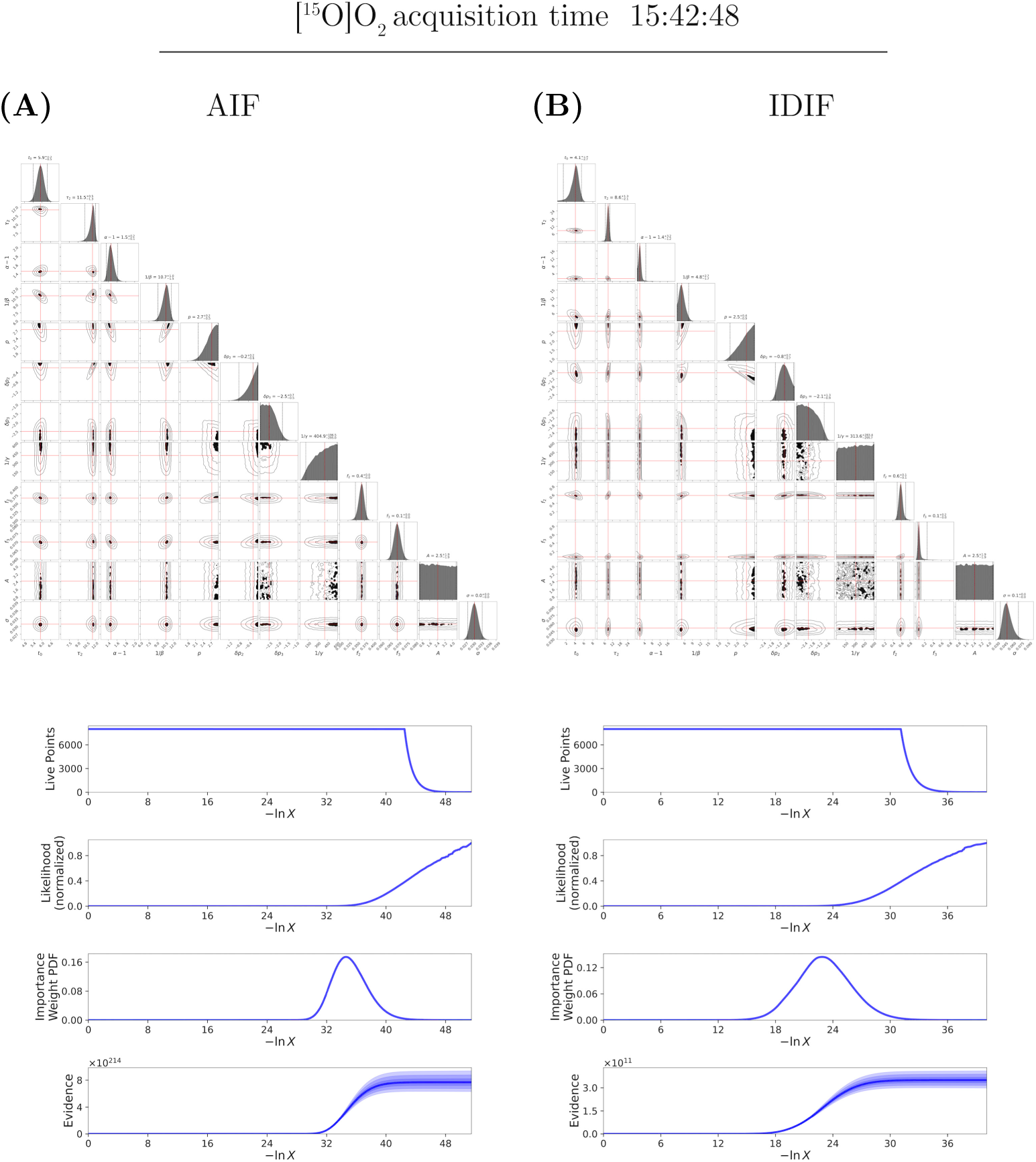
Dynamic nested sampling: parameter covariances and run plots for input functions of inhaled [^15^O]O_2_. Exemplar results for the **(A)** AIF and **(B)** IDIF of the secondary dosing of [^15^O]O_2_. **Top panels** are “corner” plots depicting covariances of model parameters. Best model parameters are indicated by red lines. **Bottom panels** show run plots. In *run plots*, nested sampling is dynamic, running to estimate the posterior as the prior configuration volume enclosing prior mass, *X*, shrinks. *Live points* are added to a fixed number of inference samples. The likelihood limit *L/L*_max_ adjusts to the shrinking prior volume *X*(*L*). The *importance weighting probability density p*(*X*) for large *dX* and small *L*(*X*) contributes to the evidence, while matched *dX* and *L*(*X*) contributes to the posterior mass and parameter estimation. The *evidence Z*(*X*) increases with posterior mass, with shades of blue indicating 1*σ*–3*σ* errors. Reference [28] supplies comprehensive computational details.

**Figure 5.**
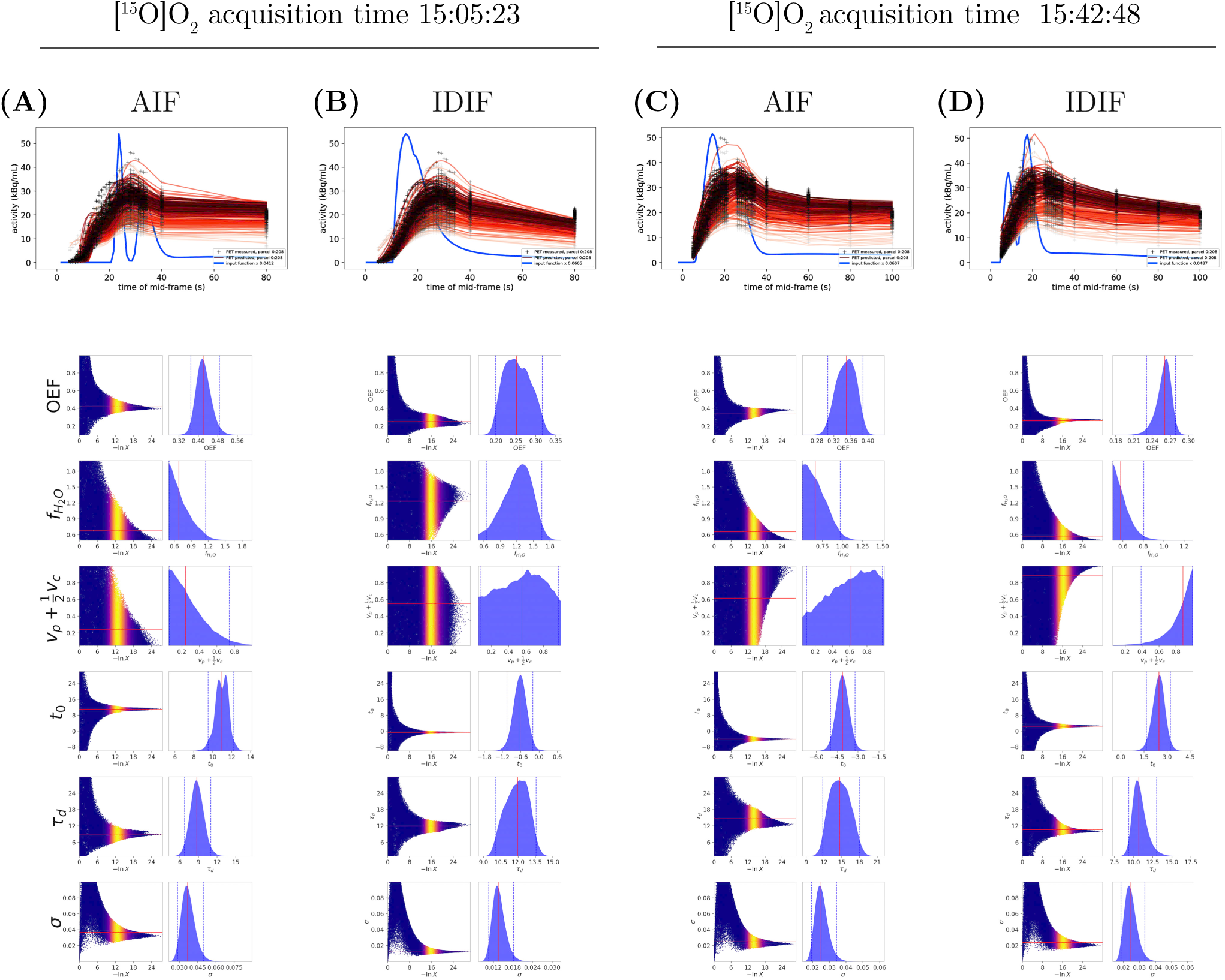
Dynamic nested sampling fits tissue time activity curves from repeated measurements of inhaled [^15^O]O_2_. As previously, columns **(A–D)** illustrate two administrations of [^15^O]O_2_, separated by 37 minutes, each observed by AIFs, **(A, C)** and IDIFs, **(B, D)**. **Topmost panels** are *residual plots*. Blue curves identically recapitulate the parametric model of unseen input functions, AIF or IDIF, previously shown in figure 3. Red curves illustrate predictions of time activity curves obtained from AIFs or IDIFs and the original model of PET for oxygen utilization [9]. Saturation and hue of red curve colors represent measured time activity curves from 109 subcortical regions from FreeSurfer (lesser saturation and lighter hue) and 100 left-sided cortical parcellations from the Schaefer atlas(greater saturation and darker hue). Crosses represent the measured time activity curves obtained from listmode with motion correction that adjusts attenuation maps with landmarks of motion. Some time points could not complete reconstruction with listmode reconstruction because of insufficient events during measurement windows. *Trace plots* indicate model parameters for tissue time activities. **Lower panels** are *trace plots* for model parameters. Compared to equation 7, *f*_H2 O_ = *µ*_metab_, *t*_0_ is the adjusted time of input function inflow which requires significant adjustment for AIFs, and *τ_d_* is a time-constant for corrective dispersion of input functions. A corrective dispersion accounted for variations of cannulation materials and methods that could not be directly measured at the time of measuring AIFs. For consistency of the bolus model, corrective dispersion was also permitted for IDIFs.

**Figure 6.**
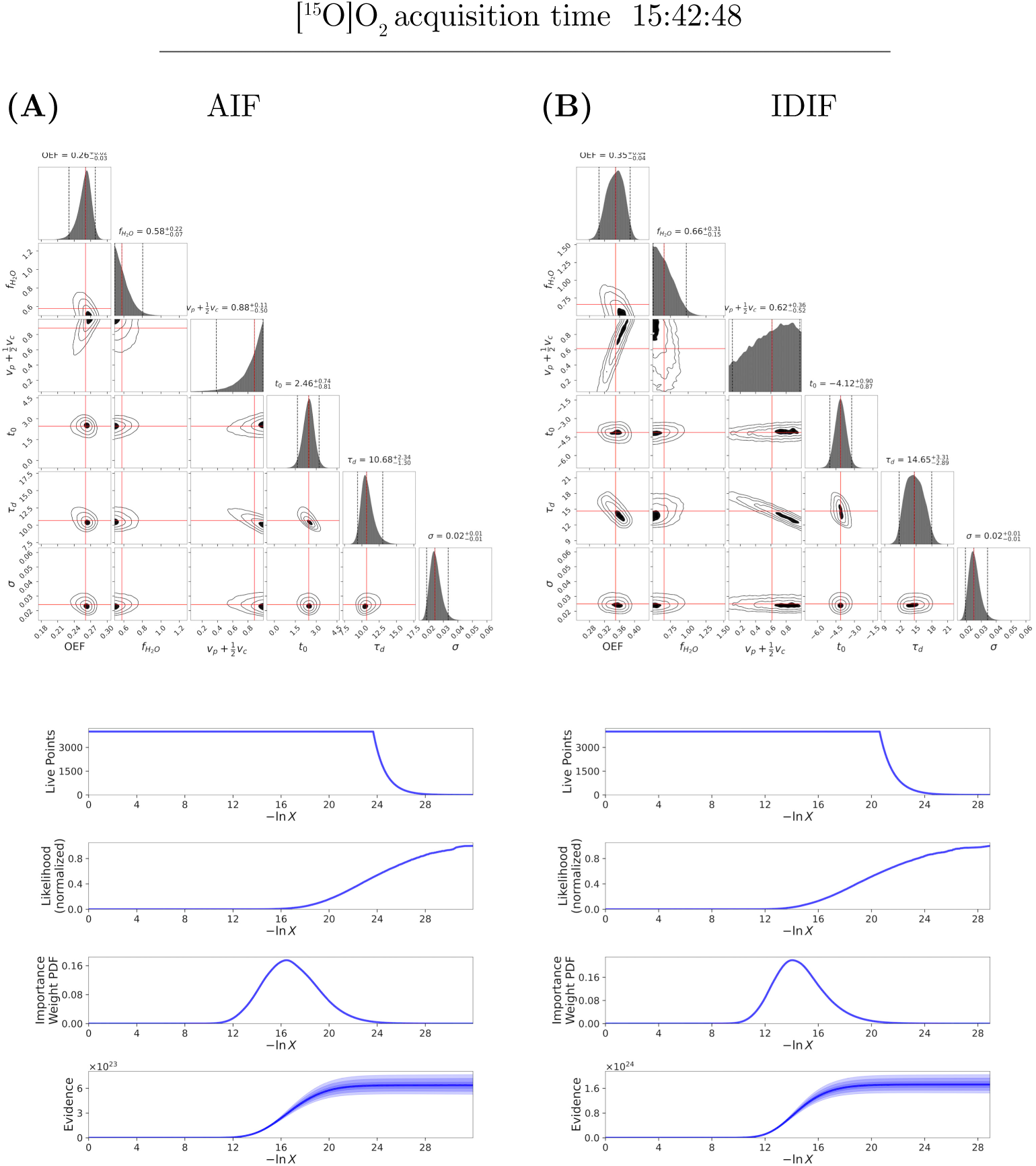
Dynamic nested sampling: parameter covariances and run plots for tissue time activity curves following inhaled [^15^O]O_2_. Exemplar results for time activity curves derived from the **(A)** AIF and **(B)** IDIF of the secondary dosing of [^15^O]O_2_. **Top panels** illustrate “corner” plots and **Bottom panels** show run plots in correspondence with figure 5, and as described for figure 4.

### AIFs have significant delays and dispersions

Arterial blood measurements encompassed extraction of blood from the radial artery cannulation, through a rigid-wall wide-bore arterial catheter, two stopcocks, a narrow-bore extension catheter feeding the arterial emissions detectors, the peristaltic pump operating at 5.00 mL/min, to the final blood collection container. Antecedent catheter line calibration studies (supplemental appendix 2) estimated mean catheter transit time to be 9.9 s for blood hematocrits in normal ranges and ambient temperature of 20 *^◦^*C. The arterial blood emissions detectors were dual LYSO crystals propagating signal photons to photomultipliers and coincidence detection electronics discriminating trues within 100 ns delay. The detection board recorded counting averages at 1.000 Hz.

The high precision of coincidence counting is offset by the variable hemodynamics between the cerebral artery circulations, the radial artery circulation, and the transport through catheter lines. In the presence of tracer-bolus recirculation, the arterial-venous circuits for the cerebral arteries and the radial artery have distinct recirculation times. Consequently, a cerebral input function and radial artery input function will have wave packets possessing distinct phase velocities, with mixing after the first recirculation. For bolus dispersion time constant shorter than recirculation time, separated recirculating peaks reveal the recirculation time. For bolus dispersion time constant longer than recirculation time, the apparent dispersion will depend on recirculation time. Finally, catheter lines impose additional significant delay and dispersion which obscure bolus wave packets *in vivo*. We have estimated catheter line delay and dispersion with Heaviside impulse-response calibration studies using packed red blood cells diluted in heparinized plasma. Such studies approximate the detailed flow characteristics of blood drawn from the radial artery of human study participants. We applied our measurements of catheter line delay and dispersions to models, detailed below.

### IDIFs have partial volumes and coarse-graining in time

Carotid imaging measurements comprised adaptations of the method of IDIFs with MR-defined carotid centerlines [12]. We obtained high-resolution anatomy of the internal carotid arteries from time-of-flight MRA and MP-RAGE and constructed centerlines as detailed in Methods. We then examined centerlines and results from the cervical segments of the internal carotid arteries and found them comparable to those previously reported [12]. However, we also pursued use of the petrous segments for additional benefits. Petrous segments of the internal carotid arteries have negligible displacements from the oriented volume of the brain when the head and neck reposition between MR and PET scanners. Petrous segments also facilitate spatial co-registration with CT-derived attenuation maps, which in turn have spatiotemporal co-registration with listmode emissions following use of BMC. Petrous segments are surrounded by bone, minimizing spill-in effects during the lifetime of ^15^O. Additionally, because we found that re-registration of centerlines from the cervical segments was difficult to assess in imaging of [^15^O]O_2_, we opted to use petrous segments exclusively. We sought to match the temporal resolution of carotid imaging measurements with that of arterial blood measurements, while sampling sufficient emissions from carotid centerlines. To this end, we reconstructed dynamic imaging using OP-OSEM on listmode with BMC, setting frame durations to 10 s, identical to frames implemented by Fung et al. [12] We repeated the reconstructions with starting time offsets from 1 to 9 s. Collating frames by mid-frame times, we obtained 1 Hz samples of carotid centerlines using 10 s moving windows.

The physiological and anatomical specificity of carotid imaging measurements is offset by partial volume effects and lower temporal resolution of the carotid centerlines. Spill-out from the petrous segment of the internal carotid arteries was expected, but spill-in appeared negligible. Our centerline constructions had effective diameters of 1.65 mm or less. Moving-window averages for time series data has known benefits for enhancing changes of state such as occurs in bolus passage of tracer. Nevertheless, distortions of time series need characterization, which we have pursued with models, detailed below.

Figure 1 provides comprehensive views of carotid imaging measurements for administration of [^15^O]CO, [^15^O]O_2_, and [^15^O]H_2_O. For comparability with arterial blood measurements, decay corrections were not applied. As for arterial blood measurements, the origin of the time axis denotes the start of emissions scanning. Since carotid imaging measurements have frame durations of 10 s, the first emissions recording denotes the mid-frame time, which is 5 s. The gaseous tracers demonstrate immediate bolus inflow, indicating lung to carotid artery transit times near 5 s. The bolus inflow of [^15^O]H_2_O occurs at 10 – 20 s, reflecting systemic venous transit times. As for arterial blood measurements, prolonged or multi-modal bolus peaks appear for [^15^O]O_2_, but also for [^15^O]CO, supporting variable respiratory inhalation of all gaseous tracers.

Additional delay and dispersion of arterial blood measurement may mask variable respiratory inhalation, especially for SNR available to arterial blood emissions detectors.

### Phenomenology of input functions may require multiple boluses and recirculations

We accumulated data to correct catheter line delay and dispersion from AIFs. By construction, we have used moving window averaging on IDIFs. If the true but hidden tracer bolus passage in the petrous carotid artery was equivalent to the tracer bolus passage in the radial artery, a single hidden variable model could be explanatory for all observations. Figure 2 illustrates the bolus forms accommodated by our series of generalized gamma distributions. Temporal parameters *t*_0_, *τ*_2_, and *τ*_3_ describe transit times for blood circulations. Parameters for stretched exponentials are *p* and increments *δp*_2_ and *δp*_3_ which apply to recirculations. The power law exponent is *α* − 1. Time constants for exponential decay and steady-state accumulations are 1*/β* and 1*/γ*, respectively. Series coefficients *f*_2_, *f*_3_, and *f*_ss_ denote the weights of recirculating boluses. Figure 1 illustrates that the radial artery circuit appears not to be consistent with the petrous carotid artery circuit. The petrous carotid artery circuit has less dispersion, consistent with shorter circuit transit, especially through slow flowing venous paths. The total area emissions activity over time for IDIFs is lower than that for AIFs, suggesting the influence of spill-out in IDIFs. Notably, despite spill-out effects, it is the carotid artery circuit that is physiologically relevant for interpreting the kinetics of tracer uptake by the brain.

### An intravascular tracer provides precision recovery coefficients

Numerical integration of all input functions from figure 7 count annihilation events over increasing time windows per unit volume from tomography or arterial blood detectors. The scatter plots and unbiased residual plots provide graphical validations of measurement methods as advocated by Bland and Altman [29]. The quotient of IDIF over AIF annihilation counts denotes the recovery coefficient. All tracers supported recovery coefficients between 0.4 – 0.7. [^15^O]O_2_ exhibited high variability between study participants and repeat scanning for a participant, and participant 1 demonstrated high discrepancies in early portions of bolus passage between AIF and IDIF and between repeat scans. [^15^O]H_2_O exhibited smoother convergence to a recovery coefficient, but recovery coefficient varied between participants, reproducing observations by Fung and Carson [12]. The most reproducible recovery coefficients emerged for [^15^O]CO, suggesting that recovery coefficient variability arises significantly from diffusion and uptake of tracers by vascular endothelia and target tissues. Intravascular tracers are necessarily more specific for quantification of vascular input functions.

**Figure 7.**
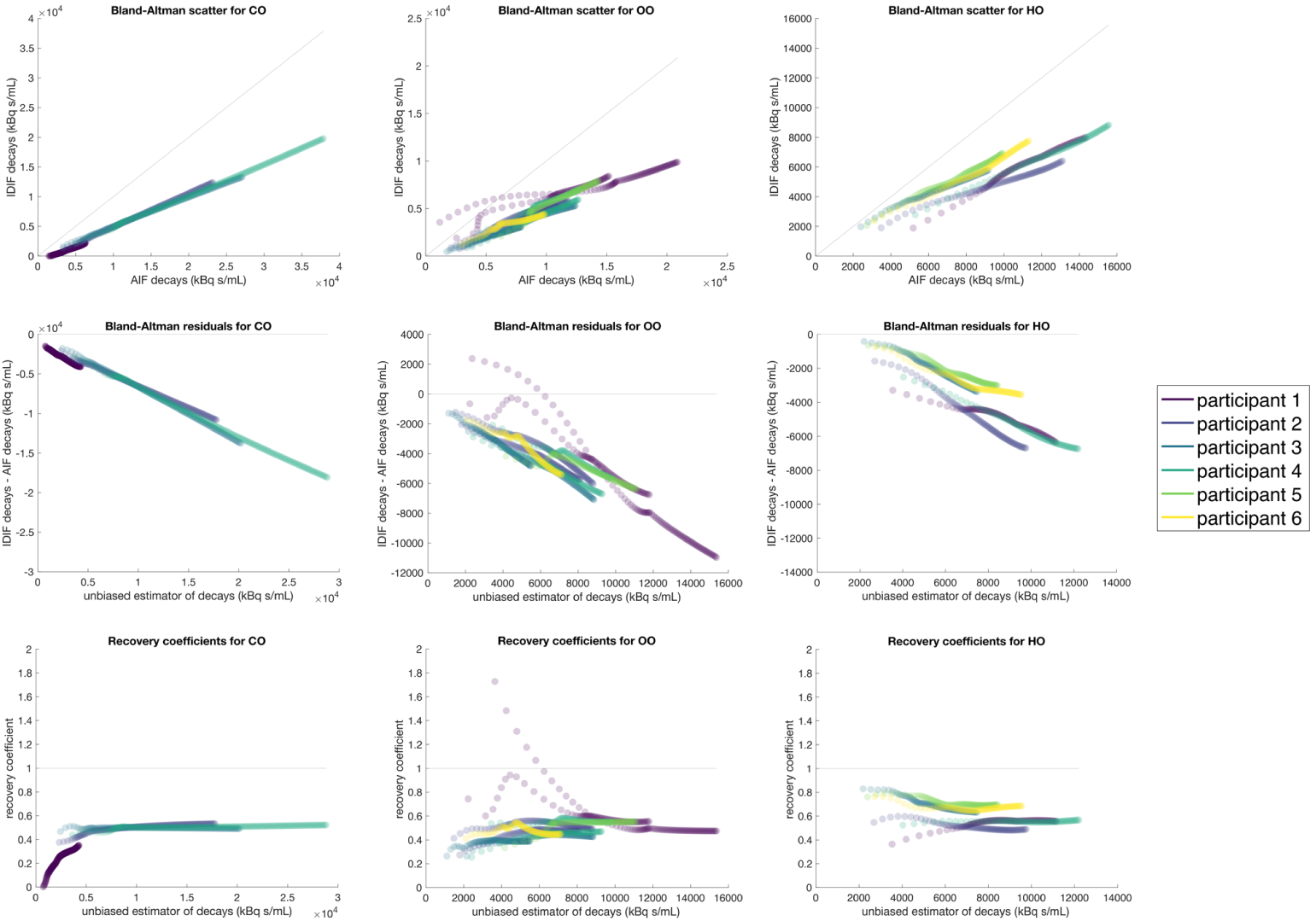
Bland-Altman graphical analyses of IDIFs and AIFs, and recovery coefficients. **Top row** illustrates the scatter plots of activity densities, expressed as cumulative decay events per unit volume for IDIFs, 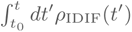, and for AIFs, 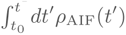. **Middle row** illustrates the residuals of cumulative decays for IDIFs compared to AIFs, plotted against the unbiased estimator, which is the mean of 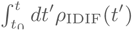 and 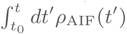. **Bottom row** illustrates the recovery coefficient, defined by the quotient 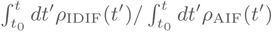 Recovery coefficient estimates were more convergent for [^15^O]CO than for [^15^O]O_2_ and [^15^O]H_2_O, possibly because of high affinity of [^15^O]CO for hemoglobins in red blood cells, making the tracer highly intravascular. The volume of distribution of [^15^O]CO matches the blood volume, the theoretical minimum for tracer kinetics. The evolution of activity densities to decay-corrected steady-state is finite compared to measurement times, but [^15^O]CO exhibits less variability than other [^15^O] tracers.

### Biological variability exceeds differences from input function

The interpretation of oxygen utilization from [15]O tracer studies arises from physiological metrics, especially CBV, CBF, OEF, and CMRO2. Several intermediary metrics that were historically imputed, *λ*, PS, *V*_post,cap_, and *f*_water_, we have inferred using nested sampling. Historical imputations are more challenging to use in modern PET that emphasizes region localization. Raincloud plots in figures 8, 10, and 12 illustrate the variability of physiological metrics across study participants, repeat scanning, and regions defined by the Schaefer 200 augmented by FreeSurfer white matter segmentations. Distributions of physiological metrics obtained from IDIFs exhibited more central tendency across persons with fewer outliers. Specifically, median values of metric for persons had narrower bounds for IDIFs compared to AIFs. The most anomalous values for physiological metrics included low median CBF with AIF, low median *V*_post,cap_ with AIF, high median OEF with AIF, and low median CMRO2 with AIF.

**Figure 8.**
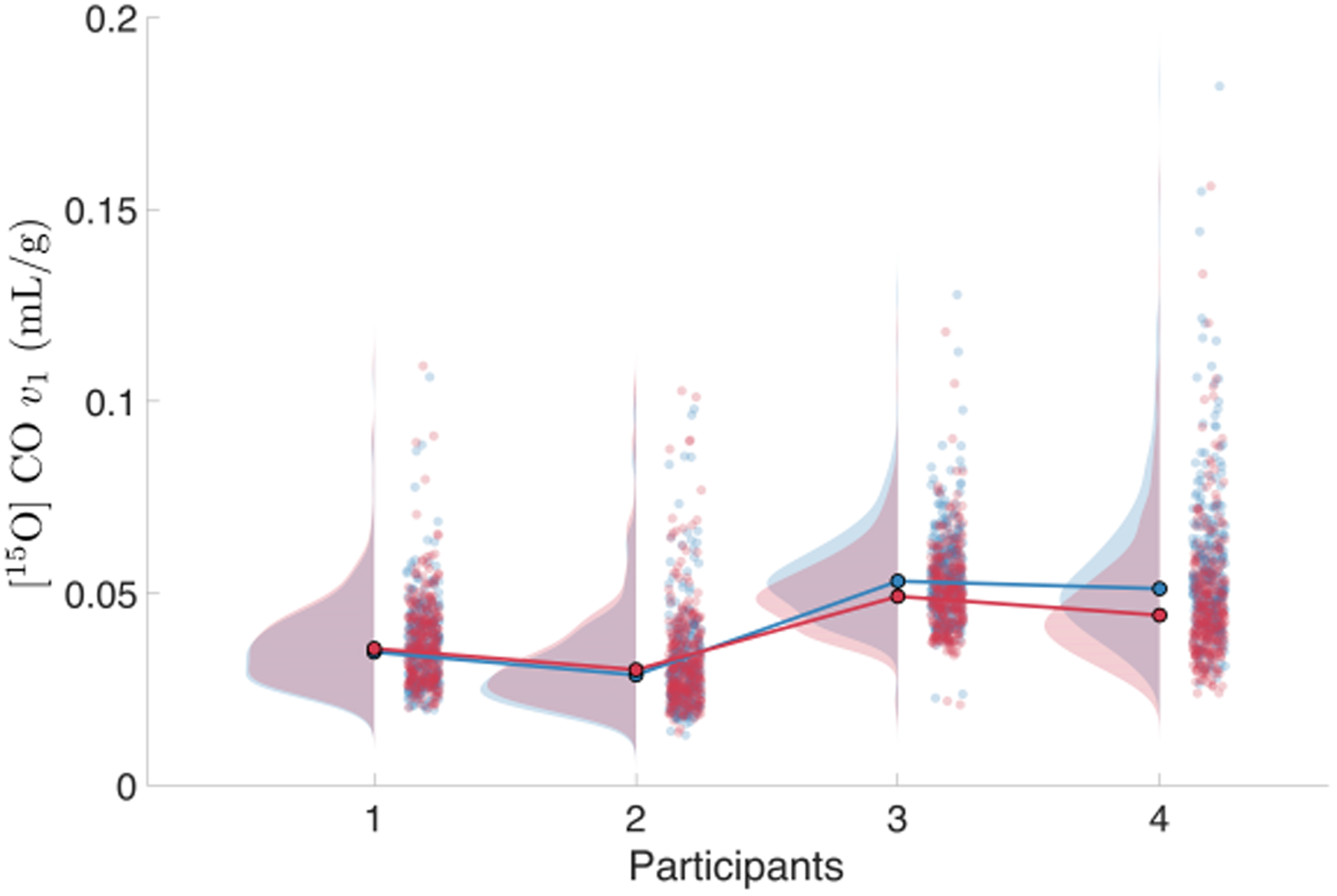
Raincloud plots demonstrate scalability of blood volume. Blue data points indicate blood volumes estimated from IDIFs sampled from the petrous carotid arteries and brain segmentations from FreeSurfer and Schaefer cortical atlases. IDIF results include rescaling by a recovery coefficient expected to depend upon characteristics of input function definitions, scanner performance, and radionuclide free paths, but may be invariant to human subjects and radiotracers. Red data points indicate blood volumes estimated from AIFs and identical brain segmentations. The similarity of histograms of volumetric samples supports the validity of the rescaling.

### Regional physiological maps have similar topographies between input functions

Figures 9, 11, and 13 illustrate the topographies of physiological metrics, on the cortical surface and selective views of subcortical structures. Each figure depicts the median value of a metric averaged over persons, to emphasize regional features. Metrics using IDIFs include rescaling by the estimated recovery coefficient. Metrics using AIFs have no rescaling. [^15^O]CO PET provides uniquely specific measurements of cerebral blood volume by virtue of its high binding affinity for hemoglobin in red blood cells. Blood volume per person is the quotient

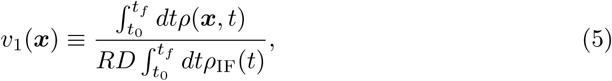

for activity density *ρ*(***x***) obtained from a Schaefer parcellation of brain tissue at location ***x*** and *ρ*_IF_ obtained from an input function, either IDIF or AIF. Early studies revealed that *v*_1_ requires approximately 120 s to attain steady state [30]. Consequently, the limits of integration are *t*_0_ = 120*s* and *t_f_* set to the end of usable time-series data.

**Figure 9.**
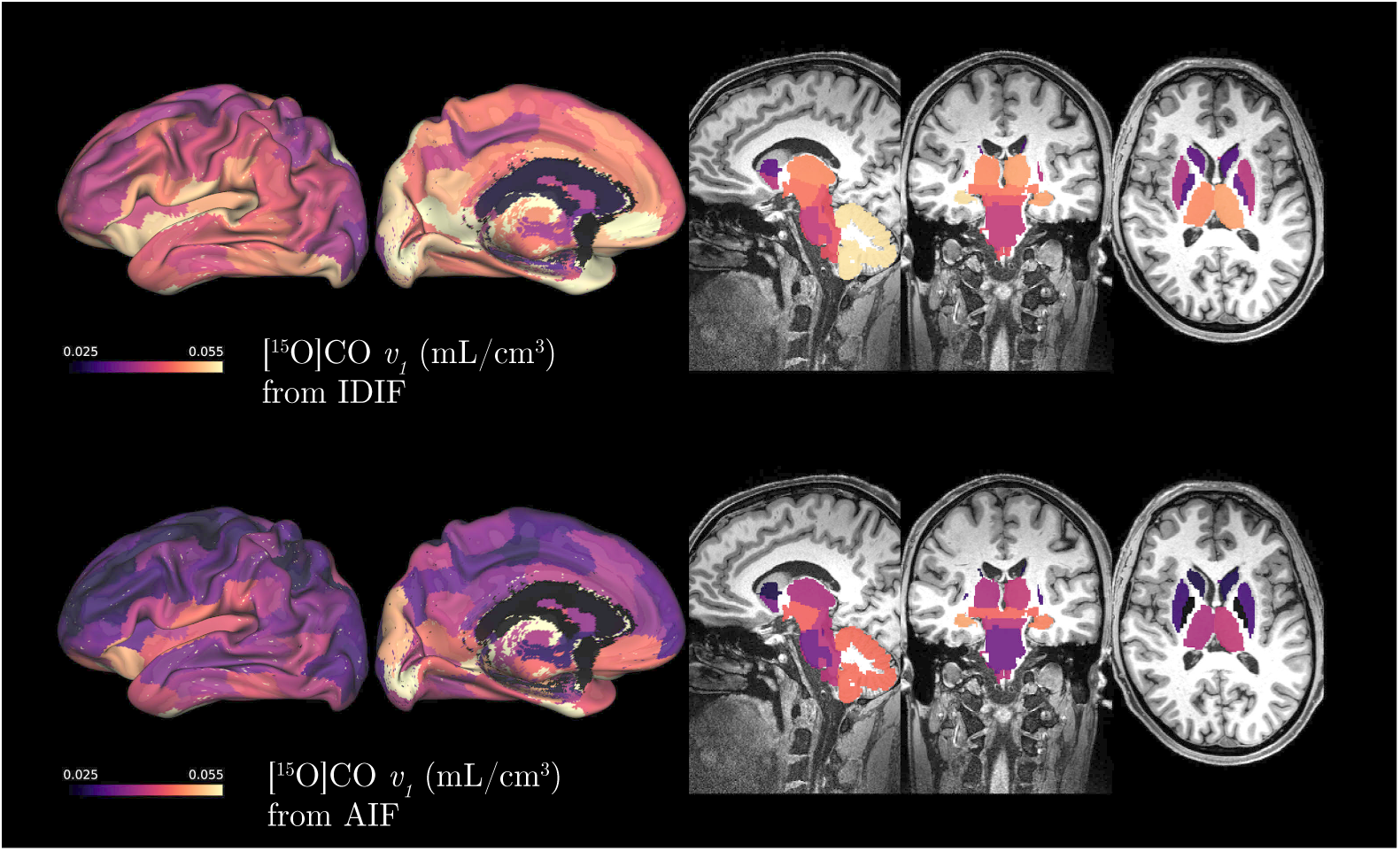
Surface and volume renderings confirm scalability of blood volume. All renderings display 200 parcellations of the Schaefer atlas and 109 subcortical segmentations from FreeSurfer. Each region indicates the median of the physiological metric over all participants. Left-sided panels show model inferences with IDIFs. Right-sided panels show inferences with AIFs. IDIFs have corrections for moving window averages, and AIFs have corrections for delays, and dispersions, as described in methods and results. Steady state conditions required for estimations of blood volumes exclude bolus first-passage in early time-frames, affecting the magnitude of blood volume maps. Topographical organization, however, remains consistent between IDIFs and AIFs.

**Figure 10.**
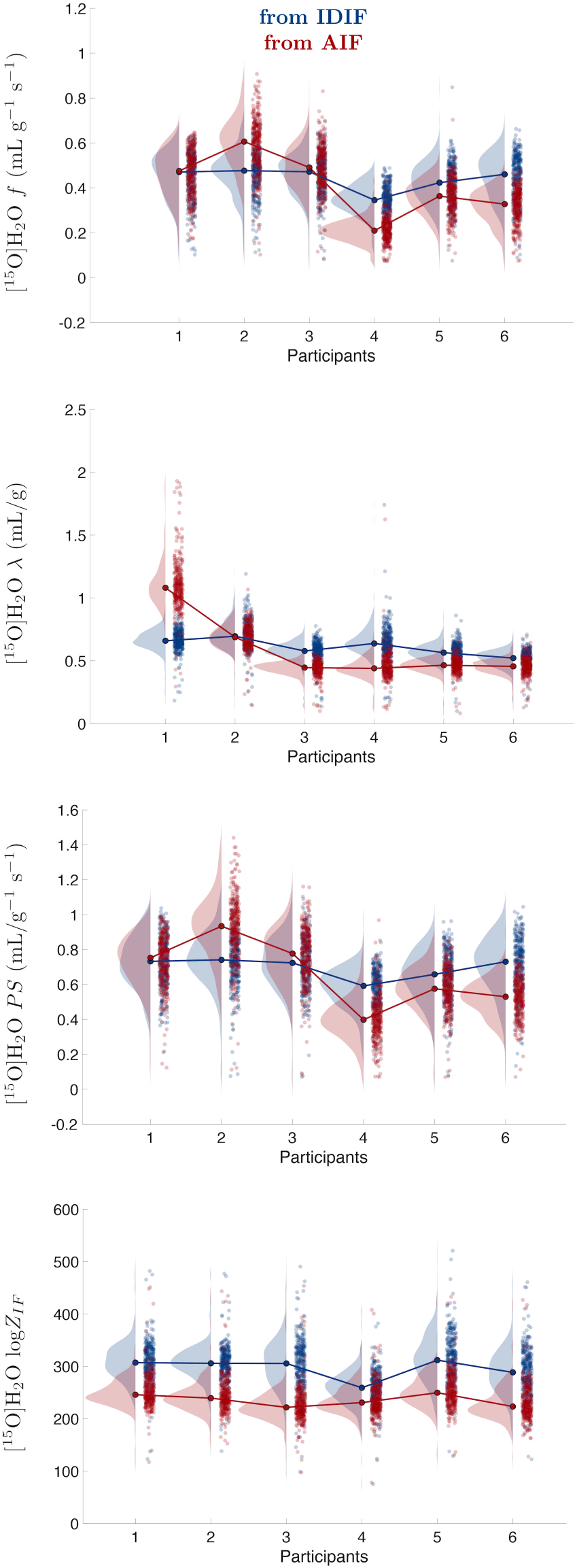
Raincloud plots for metrics of [^15^O]H_2_O. All parameter estimates fell well inside prior ranges: 0.0011 – 0.0171 mL g*^−^*^1^ s*^−^*^1^ for *f*, 0.05 – 2.00 mL/g for *λ*, and 0.0011 – 0.0283 mL g*^−^*^1^ s*^−^*^1^ for *PS*. We selected prior ranges wider than historically reported values. AIFs demonstrated metric values with more variability than metrics from IDIFs. AIF metrics were also demonstrably multimodal for *λ* and *PS*. Data evidence, log*Z*_IF_, slightly favored IDIFs.

**Figure 11.**
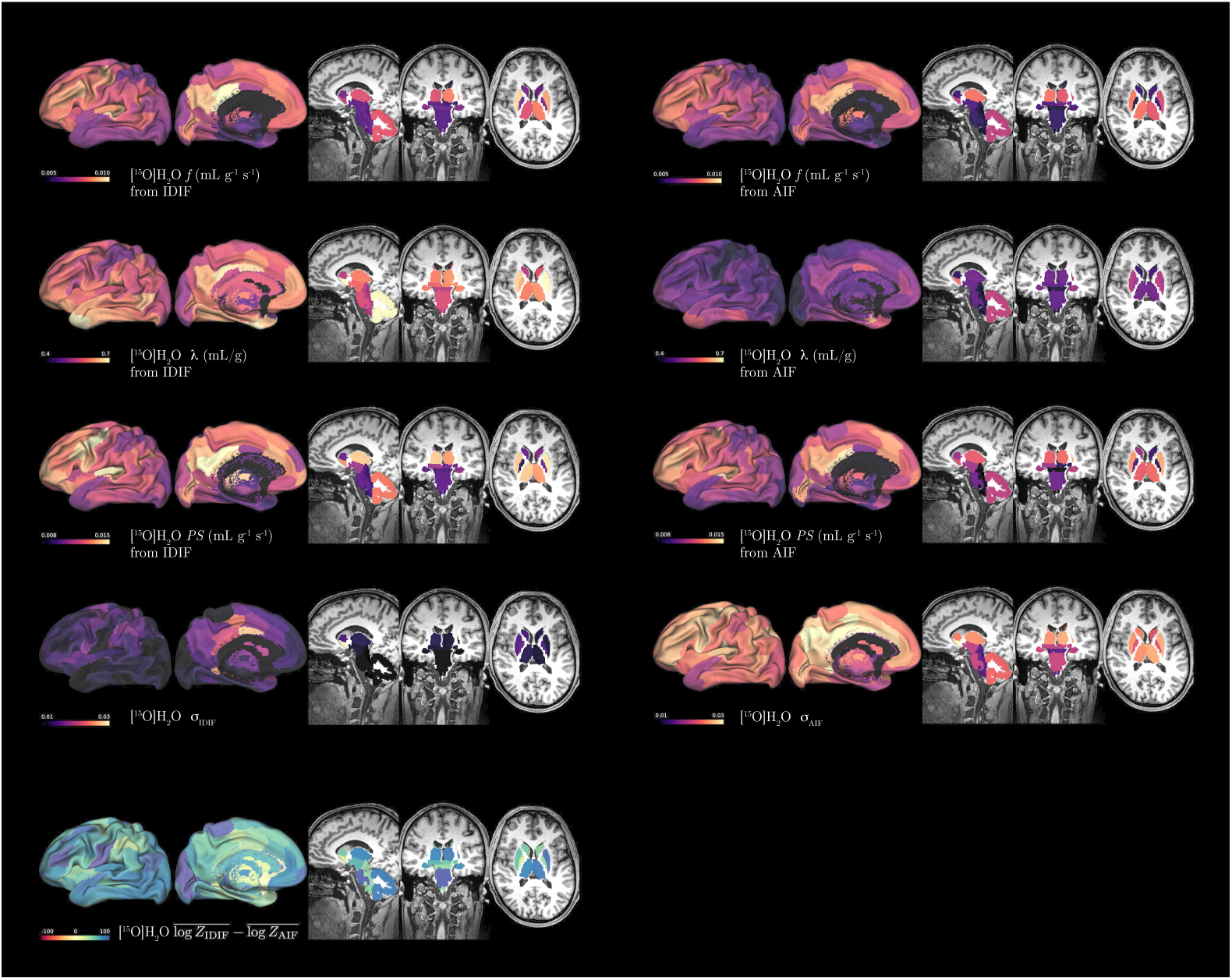
Surface and volume renderings for metrics of [^15^O]H_2_O. Flow, *f*, partition coefficient, *λ*, and permeability surface-area product, *PS*, parameterize the model of [^15^O]H_2_O emissions from tissue time activity curves and input functions. *σ* denotes the magnitude of Gaussian noise accommodating the difference between the model of [^15^O]H_2_O emissions and data. Bottom renderings illustrate data evidence favoring IDIFs over AIFs, expressed as Bayes factor log *Z*_IDIF_ *−* log *Z*_AIF_.

**Figure 12.**
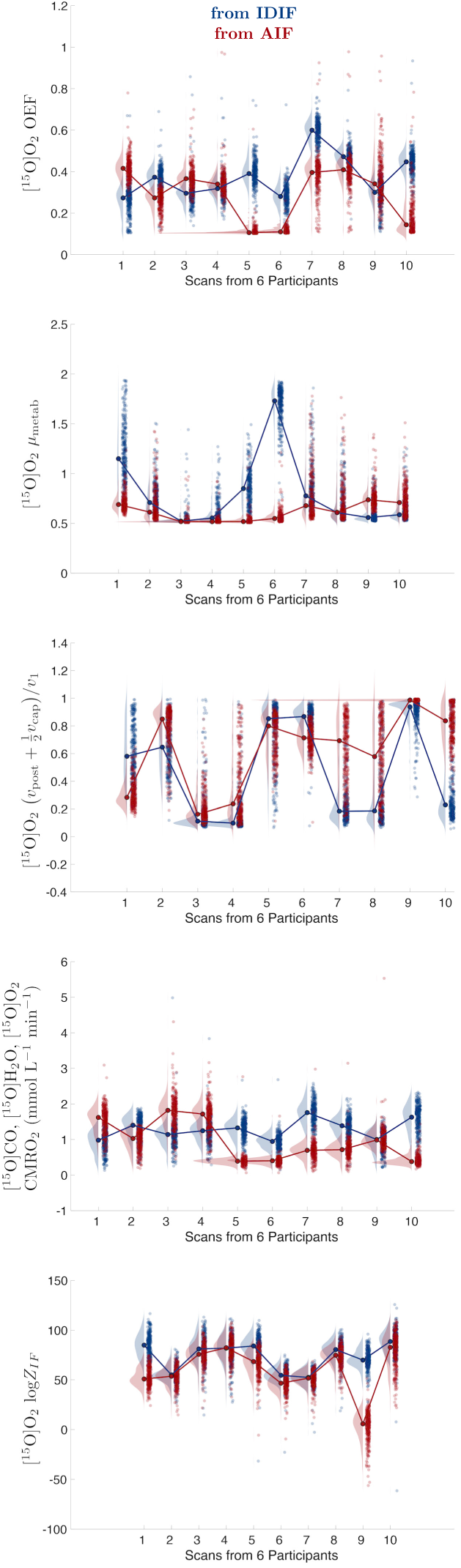
Raincloud plots for metrics of [^15^O]O_2_. Parameter prior ranges were: 0.2 – 0.8 for OEF; 0.1 – 1.9 for *µ*_metab_, the fraction of water of metabolism at 90 s; and 0.25 – 1.00 for (*v*_post_ + ^1^ *v*_cap_)*/v*_1_, the fractional post-capillary blood volume combined with fractional capillary blood volume. AIF estimates for OEF and (*v*_post_ + ^1^ *v*_cap_)*/v*_1_ frequently reached the lower prior bound. CMRO_2_ is a derived metric and AIF estimates for it were repeatedly lower than corresponding IDIF estimates. Scans 1-2 belong to participant 1. Scans 3-4 belong to participant 2. Scans 5-6 belong to participant 3. Scans 7-8 belong to participant 4. Scan 9 belongs to participant 5. Scan 10 belongs to participant 6. Notably, metrics by AIF for participant 5 had poor test-retest reproducibility. Data evidence, log*Z*_IF_, favored IDIFs.

**Figure 13.**
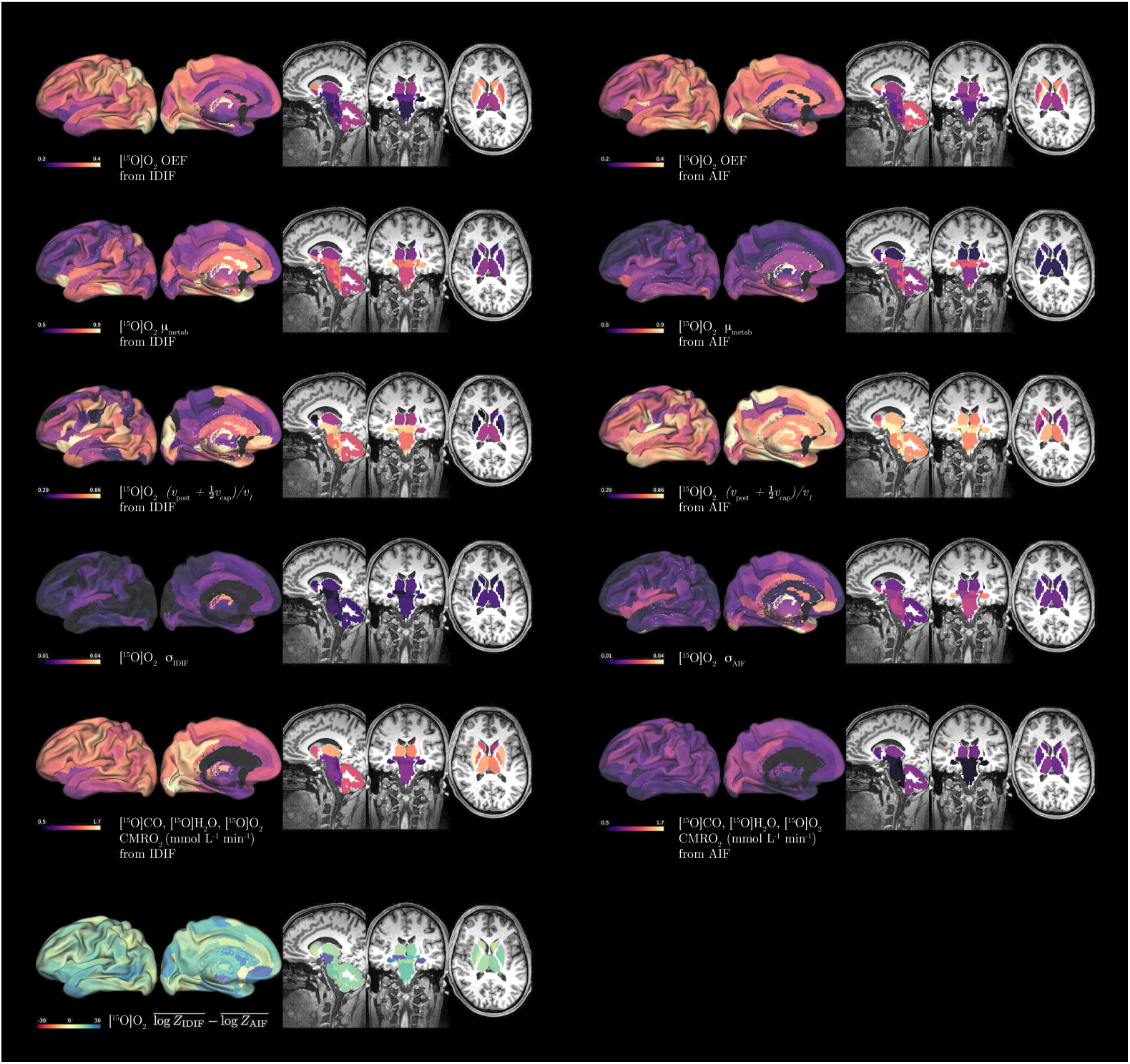
Surface and volume renderings for metrics of [^15^O]O_2_. Oxygen extraction fraction, OEF, fractional water of metabolism, *µ*_metab_, and fractional post-capillary with capillary blood volume, (*v*_post_ + ^1^ *v*_cap_)*/v*_1_, parameterize the model of [^15^O]O_2_ emissions from tissue time activity curves and input functions. *σ* denotes the magnitude of Gaussian noise accommodating the difference between the model of [^15^O]O_2_ emissions and data. Bottom renderings illustrate data evidence favoring IDIFs over AIFs, expressed as Bayes factor log*Z*_IDIF_*/Z*_AIF_.

*R* = 0.85 provides empirical adjustment of blood hematocrit for smaller vessels in brain tissue. *D* = 1.05 g/mL is the density of brain tissue. Figure 9 illustrates median person values of *v*_1_, constructed from both IDIF and AIF. *ρ*_IDIF_ contains averages from both IDIFs and AIFs by way of the recovery coefficient, *c_R_*, a mean-field quantity that also samples integrated activity densities until convergence. If integration limits for *c_R_*and for *v*_1_ were identical and identical metrics for central tendency used, then left and right panels of Figure 9 would be identical. Small quantitative contrasts reveal the effects of conventions for *t*_0_ and use of medians for persons.

[^15^O]H_2_O PET remains the standard for *in vivo* measurements of cerebral blood flow. For completeness, we retained all parameterizations of blood flow that have been reported and not yet found inconsistent with observations. Many parameterizations historically have been ignored for ease of model fitting. From conservation of mass of positron-emitting tracer and the continuity equation, we articulate

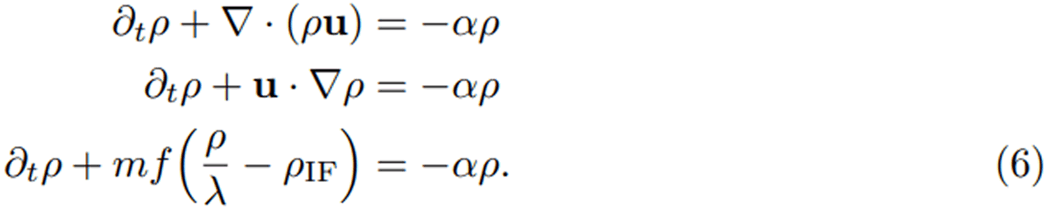

The tracer decays with constant *α* = log 2*/τ_h_*, *τ_h_*denoting the half life of [^15^O]. It is carried through incompressible blood and tissue such that the velocity field ***u*** has no divergence. Finally, we approximate the gradient by the scalar difference of tissue partitioning *ρ/λ* and input function *ρ*_IF_. The parallel velocity field includes flow *mf* with *m* = 1–*e^−P^ ^S/f^* providing empirically observed corrections using a permeability surface-area product *PS*. Figure 11 illustrates median person estimates of the variable parameters enumerated, constructed from both IDIF and AIF. IDIF provided greater estimates of flow *f* and partition coefficient *λ*, but lower estimates of *PS*. Inspecting ratio maps of IDIF over AIF did not reveal scalar proportionality of regions.

[^15^O]O_2_ PET provides the most direct measurement of oxygen metabolism *in vivo*. Its models also follow conservation of tracer mass and the continuity equation, but add fluxes for the metabolism of [^15^O]O_2_ to [^15^O]H_2_O. The metabolism may be regarded as a change of state for subpopulations of the tracer experiencing physiologic oxidative phosphorylation. Measured activity density *ρ* = *ξ* + *η* arises separately from *ξ* for [^15^O]H_2_O and *η* for [^15^O]O_2_. We articulate extensions to equations 6 to be

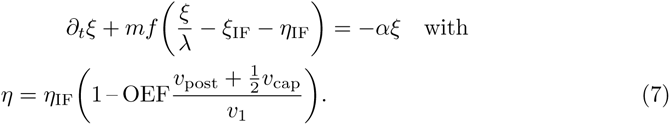

Noteworthy are the necessity of estimating *η*_IF_ from [^15^O]O_2_ and having access to measurements of only *ρ* = *ξ* + *η*. Additionally, the activity density of the water of metabolism was estimated to be, *ξ*(*t* = 90 s) ≈ 0.5 *ρ*(*t* = 90 s). We defined variable parameter *µ*_metab_ ≡ *ξ*(*t* = 90 s)*/η*(*t* = 90 s). Figure 13 illustrates median person estimates of the additional parameters introduced for [^15^O]O_2_ PET.

### Nested sampling Bayes factors favor IDIFs

The expected logarithms of evidence, log *Z*_IDIF_ and log *Z*_AIF_), provide comparative Bayes factors for model selection consistent with data. The Bayes factors illustrated for scans of [^15^O]H_2_O and [^15^O]O_2_ in the final panels of figures 11 and 13 favor IDIFs over AIFs for all regions of the brain. Furthermore, our dynamic nested sampling implementations estimated Gaussian noise sources, *σ*_IDIF_ and *σ*_AIF_ to accommodate mismatches of models and data. Noise estimates illustrated in the fourth rows of figures 11 and 13 also favor IDIFs over AIFs. We clarify that the inferences of primary interest are the estimation of physiologic kinetic metrics such as blood flow or metabolic rates, and our concluding model selections concern these. The local physiology has been directly observed by dynamic PET tomography. We have also made intermediate inferences of the waveform of model input functions which are nonlocal, by convention, and have been indirectly observed by IDIFs and AIFs, the former involving instrumental measurements that share significant mutual information with tomography, but the latter involving invasive procedural measurements that possess relevant external information. We have consistently inferred physiology from model input functions.

Model input functions thereby distinguish IDIFs and AIFs only by the parameterization of bolus model equation 1, minimizing confounding effects of instrumental information. The inference methods for physiology and input functions have also been implemented with high emphasis for consistency of numerical implementation details.

## Discussion

Two of the major goals in methodological biomedical research are 1) to improve the precision and accuracy of measurements to be more applicable at the individual level, and 2) to lower methodological risk and costs such that they can be better applied to large cohorts and clinical populations. Developing an IDIF method to replace invasive AIF measurements for quantitative brain PET imaging accomplishes both goals. Here we took advantage of improvements in modern PET scanner technology and inferential modeling to calculate IDIFs for ^15^O metabolic PET and find that they are at least comparable to AIFs, and potentially superior [31].

Modern choices for precision data inference include supervised learning, which builds upon *a priori* models, and unsupervised learning, which seeks out structural relationships between data with objectives such as data-driven dimensionality reduction. This work considers supervised learning in the context of Bayesian inference. Bayesian supervised learning is more capable than ever, and parameter estimations with model evidence are especially transparent with nested sampling [32], [33].

The parametric models for tracer kinetics of oxygen utilization that we have presented bring the biases of models that have been historically corroborated by meticulous measurements [9], [22], [25], [30]. We have taken the view of over-completeness in our model descriptions, given the enhanced capability of our dynamic nested sampling implementations for discovering subsets of the model parameter space that are most relevant for data [28]. Models presented in this work incorporated both the underlying physical and physiologic processes as well as detailed measurement procedures. This specificity allows generalizations regarding input functions for kinetics which, after corrections of measurement effects, generalize as input functions for populations [34].

However, we found that waveforms for AIFs and IDIFs are not mutually consistent after corrections of measurement effects such as dispersion, delay, or effects of moving windows for imaging frames. This is partly a matter of temporal resolution, but the role of inconsistent circulation circuits also appears evident. In retrospect, the simplifying assumption of ignoring recirculation transport was historically sensible, but as the needs for precision measurements grow, these assumptions draw reconsideration. Modern scanner instrumentation with long bore detection of emissions promises ready sampling of cardiac or aortic IDIFs. Nevertheless, even these robust measures of IDIFs may have measurable inconsistencies with IDIFs sampling the internal carotid arteries.

Notable considerations involve inaccuracies arising from the measurement process for AIFs. Ex vivo arterial blood sampling has substantively greater measurement degrees of freedom compared to emissions counting of arterial blood from imaging. Arterial cannulation involves extraction of blood through approximately 50 cm of nonbiological catheter line materials, without physiological pulsatility, with altered blood pressures, and temperature gradients typically ≈ 1.7 *^◦^*C. These induce significant variations of spectral modes of flow, viscosity, dispersion and delay of time-series measures. These conditions also promote clotting, especially with repeated starting and stopping of arterial blood sampling as required for studies of oxygen utilization. Variability of blood transport during prolonged arterial blood sampling is likely dynamic, but no available experimental metrics can track the dynamic variations of blood as fluid. Practical considerations also abound. Historically, heparinization of catheter lines has been used to minimize clotting effects, but modern best practices for invasive clinical research make heparinization challenging in the face of very small but devastating risks of future heparin-induced thrombocytopenia [35].

Traditional models have benefits deriving from conservation principles such as the continuity equation of fluid dynamics [36]. First passage likely suffices for first-order description of the kinetics of delivery, described by parameters of flow. However, parameters of second-order transport such as the partition function between blood and tissue, *λ* [26], and extraction fractions, including *m* = 1– exp(−*PS/f*) [25] and OEF [9], are more meaningful for regional physiological inferences. OEF is essential for estimating oxygen metabolic rates. Inconsistent descriptions of recirculations between input functions and tissue time activity curves are likely more consequential for these metrics. This would explain, in part, the favorable Bayesian data evidence of IDIFs over AIFs.

We found the estimated recovery coefficient to be most consistent and informative for intravascular [^15^O]CO. All radiotracers exhibit transient pharmacokinetics and [^15^O]CO is known to exhibits transients for at least 120 s. Nevertheless, compared to other [^15^O] tracers, [^15^O]CO appears mostly likely to generalize as multiple study participants demonstrate consistent recovery coefficient for [^15^O]CO. Conceivably, the generalizability of recovery coefficient for [^15^O]CO may arise from its highly specific volume of distribution, binding to hemoglobin of red blood cells with *>*98 % avidity in our measurements. In contrast, [^15^O]H_2_O and [^15^O]O_2_ readily diffuse into tissues and the oxygen is readily metabolized. Processes of diffusion and metabolism add substantial transients which can persist through many half-lives of [^15^O].

As a practical clinical matter, the importance of the radial artery over the femoral artery has increased greatly for modern interventional procedures involving arterial access in the radiological and cardiology settings. While overt complications associated with radial artery manipulations in clinical settings are very low, ≈ 0.1%, repeated radial artery manipulation increases the risk of radial artery occlusion, including subclinical cases that might only be evident in the future when radial artery catherization is required. Thus, the development and use of methods that avoid radial artery manipulation are increasingly warranted, particularly in clinical populations. Use of IDIFs should accordingly now be assessed alongside the limitations and risks of AIFs [37].

### Limitations

Study size was limited by risk-benefits for study participants in invasive scan sessions with up to four scans of [^15^O] tracers. The small number of participants, however, facilitated detailed examination of model features and their performance for explaining data, and a growing emphasis on precision imaging that has been fruitful in recent studies [38], [39]. Bayesian methods such as dynamic nested sampling are especially suited for precision examination of small numbers of participants [32]. Notably, our reported results are less applicable to models that demand measured metabolites for every scan, as is the case for receptor binding studies [34].

## Conclusions

For research programs which can manage greater computational resources, modern Bayesian methods, such as dynamic nested sampling, can provide exceptionally precise inferences on data and models. For [^15^O] tracers, we found Bayesian data evidence to be higher for IDIFs over AIFs when using historically validated models of hemodynamics, oxygen metabolism, and tracer bolus transport. We regard our results in light of the operational challenges of invasive, serial arterial blood measurements in the setting of [^15^O] tracers, especially inhaled [^15^O] O_2_. Historically, similarly invasive studies of [^15^O] relied on accessible scanner bores that allowed for short-length catheters that minimized dispersions, higher dosing for greater signal-to-noise, reduced scattering artifacts by collimated 2D detection modules, systemic heparinization to avoid clotting during repeated series of arterial sampling, and motivated study participants who would tolerate challenging study procedures. These historical protocols are no longer compatible with modern PET scanners and the necessity of highly reproducible human studies that can scale to large numbers of study participants. The primary liability of IDIFs, apart from high computational demands, appears to be partial volume effects, for which empirical construction of scalar recovery coefficients have nevertheless yielded promising Bayesian data evidence.

## Acknowledgments

We are particularly grateful to our research participants for their altruism. We thank Tony A. Durbin, Jennifer Byers, Kim Casey, and Hussain Jafri for their assistance with human imaging and laboratory work. We acknowledge the directors and staff of the Neuroimaging Labs Research Center, Knight Alzheimer’s Disease Research Center, Center for Clinical Imaging Research (CCIR), and the Washington University cyclotron facility for making this research possible. We gratefully acknowledge research funding from NIH R01AG053503, R01AG057536, RF1AG073210, RF1AG074992, R21AG085245, the Mallinckrodt Institute of Radiology, and the McDonnell Foundation for Systems Neuroscience at Washington University.

## Declarations

JSS is a consultant for Sora Neurosciences. JJL is a consultant for Eisai Co., Ltd.

